# Dependence of perceptual saccadic suppression on peri-saccadic image flow properties and luminance contrast polarity

**DOI:** 10.1101/2020.11.26.399840

**Authors:** Matthias P. Baumann, Saad Idrees, Thomas A Münch, Ziad M. Hafed

**Affiliations:** Werner Reichardt Centre for Integrative Neuroscience, Tübingen University, Tübingen, Germany; Hertie Institute for Clinical Brain Research, Tübingen University, Tübingen, Germany

**Keywords:** Saccadic suppression, stimulus polarity, luminance contrast, trans-saccadic perception, ON channels, OFF channels, retina

## Abstract

Across saccades, perceptual detectability of brief visual stimuli is strongly diminished. We recently observed that this perceptual suppression phenomenon is jumpstarted in the retina, suggesting that the phenomenon might be significantly more visual in nature than normally acknowledged. Here, we explicitly compared saccadic suppression strength when saccades were made across a uniform image of constant luminance versus when saccades were made across image patches of different luminance, width, and trans-saccadic luminance polarity. We measured perceptual contrast thresholds of human subjects for brief peri-saccadic flashes of positive (luminance increments) or negative (luminance decrements) polarity. Perceptual thresholds were >6-7 times higher when saccades translated a luminance stripe or edge across the retina than when saccades were made over a completely uniform image patch. Critically, both background luminance and flash luminance polarity relative to the background strongly modulated peri-saccadic contrast thresholds. In addition, all of these very same visual dependencies also occurred in the absence of any saccades, but with qualitatively similar rapid translations of image patches across the retina. Our results support the notion that perceptual saccadic suppression may be fundamentally a visual phenomenon, and they motivate neurophysiological and theoretical investigations on the role of saccadic eye movement commands in modulating its properties.

## Introduction

Due to their ballistic nature, visual input across saccades invariably includes periods of large image uncertainty. Perceptual cancelation during such periods takes place, resulting in a seamless subjective visual experience despite the occurrence of saccades as often as several times in just one second (Melcher, 2011; Wurtz, 2008; Wurtz, Joiner, & Berman, 2011; Zimmermann & Bremmer, 2016). In the laboratory, the properties of peri-saccadic vision have been studied by presenting very brief and fleeting “probe” stimuli around the time of saccadic eye movements. Such stimuli act as impulses that essentially capture the instantaneous and momentary state of the visual system (Matin, 1974; Ross, Morrone, Goldberg, & Burr, 2001).

When presenting very brief peri-saccadic visual stimuli, a prominent observation is a massive, but transient, suppression of visual sensitivity. In this striking phenomenon, a visual probe goes completely unnoticed if presented within approximately +/- 50 ms from saccade onset, even if it would be detected effortlessly when presented under fixation (Beeler, 1967; Brooks & Fuchs, 1975; Idrees, Baumann, Franke, Munch, & Hafed, 2020; Latour, 1962). This phenomenon was labeled saccadic suppression (Zuber & Stark, 1966), and it has attracted investigation by vision scientists for many decades (Binda & Morrone, 2018; Castet, Jeanjean, & Masson, 2001; Castet & Masson, 2000; Matin, 1974; Schweitzer & Rolfs, 2020; Wurtz, 2008). Saccadic suppression is robust, and it occurs for both reflexive and deliberate saccades (Gremmler & Lappe, 2017). It also seems to occur for saccades of all sizes, including microsaccades (Beeler, 1967; Chen & Hafed, 2017; Hafed & Krauzlis, 2010; Zuber & Stark, 1966), and it even acts to shape the long term dynamics of visual sensitivity well after the eye movements (J. Bellet, Chen, & Hafed, 2017; Benedetto & Morrone, 2017). Most interestingly, neural responses to visual probes in a variety of brain areas are also suppressed if the probes occur peri-saccadically (Berman, Cavanaugh, McAlonan, & Wurtz, 2017; Bremmer, Kubischik, Hoffmann, & Krekelberg, 2009; Chen & Hafed, 2017; Chen, Ignashchenkova, Thier, & Hafed, 2015; Hafed & Krauzlis, 2010; Idrees, Baumann, Franke, et al., 2020; Idrees, Baumann, Korympidou, et al., 2020; Kagan, Gur, & Snodderly, 2008; Reppas, Usrey, & Reid, 2002; Robinson & Wurtz, 1976).

For perhaps as long as saccadic suppression has been investigated, there have been debates on its origins. On the one hand, according to the “active hypothesis”, it has been suggested that suppression relies on knowledge of saccade generation commands to actively suppress visual sensitivity, through efference copies or corollary discharge from (pre-) motor areas (Beeler, 1967; Diamond, Ross, & Morrone, 2000; Duffy & Lombroso, 1968; Zuber & Stark, 1966). Perceptually, a showpiece for this hypothesis was the observation that saccadic suppression can be selective (Burr, Morrone, & Ross, 1994): brief flashes of low spatial frequency patterns are suppressed much more than brief flashes of high spatial frequency patterns (Burr et al., 1994). This observation was interpreted as evidence for an active movement-related signal specifically targeting the magnocellular visual pathway, which is sensitive to low spatial frequencies (DeYoe & Van Essen, 1988; Merigan & Maunsell, 1993). On the other hand, according to the “visual hypothesis”, it is the visual consequences of saccades that jumpstart suppression. This is supported by the observation that brief probe flashes near the onset of global image translations (similar to those caused by saccades) are also perceptually suppressed (Adey & Noda, 1973; Brooks & Fuchs, 1975; Brooks, Impelman, & Lum, 1981; Castet et al., 2001; Diamond et al., 2000; Idrees, Baumann, Franke, et al., 2020; Mackay, 1970; Mitrani, Mateeff, & Yakimoff, 1971; Mitrani, Radil-Weiss, Yakimoff, Mateeff, & Bozkov, 1975; Mitrani, Yakimoff, & Mateeff, 1973; Noda & Adey, 1974; Yakimoff, Mitrani, & Mateeff, 1974). Recently, we also found that even selective suppression of low spatial frequency patterns may be entirely visual in origin: both selective and unselective suppression can happen with or without saccades, simply as a function of visual context (Idrees, Baumann, Franke, et al., 2020). A key observation was that saccade-like image translations activate, rather than suppress, the retinal image processing cascade and result in afferent activity bursts in retinal ganglion cells; responses to subsequent flashes are, in turn, suppressed through retinal-circuit (and downstream) visual interactions (Idrees, Baumann, Franke, et al., 2020; Idrees, Baumann, Korympidou, et al., 2020). Thus, from a visual perspective, perceptual saccadic suppression involves visual-visual interactions between two kinds of signals: (1) the visual consequences of saccadic eyeball rotations; and (2) the visual consequences of flash onsets (Idrees, Baumann, Franke, et al., 2020).

Here, we explored these visual-visual interactions in more detail at the perceptual level. We varied the image conditions across which gaze translated during saccades, and we also tested image translations in the absence of saccades. We additionally explored influences of flash polarity (luminance increments or decrements relative to the background), motivated by quantitative differences between ON and OFF retinal pathways in saccadic suppression (Idrees, Baumann, Korympidou, et al., 2020). We found that saccadic suppression exhibits a large diversity of visual dependencies, which also emerge with image translations in the absence of saccades. Our results strongly motivate revisiting both the movement-related and visual components of saccadic suppression, and investigating how saccade movement commands may interact with visual-visual interactions in shaping trans-saccadic visual perception.

## Methods

### Subjects and ethical approvals

We collected data from 6 human subjects (3 females) who provided informed consent. Two subjects were authors of this study (MPB and SI), and the others were naïve to the purposes of the experiments. The subjects’ ages were in the range of 22-32 years, and the subjects were compensated 10 euros per session of approximately 60 minutes each. Each subject’s data were collected across 10 individual sessions, resulting in 60 sessions in total. The experiments were approved by ethics committees at the Medical Faculty of Tübingen University, and they were in accordance with the Declaration of Helsinki.

### Laboratory setup

The laboratory setup was similar to that described in recent studies from our group (Bogadhi, Buonocore, & Hafed, 2020; Idrees, Baumann, Franke, et al., 2020). Subjects sat in front of a CRT display placed 57 cm in front of their eyes, and the display had a resolution of 41 pixels/deg and a refresh rate of 85 Hz. In visual angles, the display spanned approximately 34 deg horizontally and 26 deg vertically. We stabilized subjects’ heads in front of the display using a custom-built device, consisting of a chin rest, forehead rest, temple guides and a head-band (Hafed, 2013). In addition, we tracked eye movements using a video-based eye tracker (Eyelink 1000, SR Research Ltd, Canada) that was placed on the desktop under the display and aimed at the left eye. Before running the experiments, we linearized the display (8-bit resolution), such that we had equal steps of luminance increments or decrements (relative to the background luminance) for the different experimental variants (Idrees, Baumann, Franke, et al., 2020). All experiments were controlled using the Psychophysics Toolbox (Brainard, 1997; Kleiner, Brainard, & Pelli, 2007; Pelli, 1997), which also synchronized data from the eye tracker using Eyelink extensions of the toolbox (Cornelissen, Peters, & Palmer, 2002). Data and events were stored and later analyzed using Matlab.

### Experimental procedures

Our general approach was to present a brief probe flash (1 display frame; approximately 12 ms) at a specific time relative to a visual transient. In some conditions, such a visual transient was caused by a real saccade shifting the retinal image globally. In other conditions, it was caused by either a luminance step in the display during fixation or by a transient translation of the display image (to simulate a saccade-like image displacement). The brief probe was presented at one of multiple possible display locations, which were fixed in position on the display for all experiments. Subjects had to identify, on every trial, at which of the locations the probe flash appeared. Therefore, the experiment utilized a multiple-alternative forced choice design. If subjects experienced suppression of their perceptual sensitivity, then their proportion of correct responses was expected to significantly decrease relative to normal perceptual performance.

To obtain a sensitive measure of perceptual sensitivity across the different conditions, we varied the flash contrast (luminance amplitude, either as an increment or decrement from background luminance) across all trials, and we collected full psychometric curves of perceptual detectability. This provided a much more sensitive way of assessing perceptual thresholds across conditions when compared to just plotting the percentage of correct trials for just one flash contrast (Idrees, Baumann, Franke, et al., 2020).

Across ten sessions per subject, we tested 5 different visual transient conditions, along with two different background luminance conditions at the time of perceptual discrimination (i.e. at the time of probe flash occurrence). We describe these conditions in detail next. Each condition was tested separately in two sessions per subject.

#### (1) Real saccades across a uniform background

In this condition, subjects started by fixating a white fixation spot that appeared at 11.8 deg eccentricity from display center, and located either to the right or left of display center as follows: 11.2 deg to the right and either 3.8 deg above or below display center; or 11.2 deg to the left and either 3.8 deg above or below display center (Fig. 1A, top). The fixation spot was a square of 7.3 by 7.3 min arc size and 142.8 cd/m^2^ luminance (Idrees, Baumann, Franke, et al., 2020). After a random delay of 800-1700 ms after initial fixation spot appearance, the fixation spot disappeared, and it simultaneously re-appeared at display center. This was the instruction to generate a saccade towards the new spot location (Fig. 1A, top). We then initiated an automatic program to detect saccade onset. This program was described previously (Chen & Hafed, 2013), and it was also similar to what we used recently in a similar context to the present one (Idrees, Baumann, Franke, et al., 2020). Briefly, this saccade detection program collected a recent history of eye position samples and used those to obtain a running estimate of eye speed. If such speed exceeded a threshold, a saccade was detected. Upon saccade detection, we presented a single-frame probe flash as the perceptual discrimination stimulus (Fig. 1A, top). The probe flash occurred after one of 4 possible time points after online saccade detection: 24, 35, 47, or 59 ms. These time points were guided by our experience in a similar context (Idrees, Baumann, Franke, et al., 2020), and they allowed us to obtain a time course of recovery in perceptual sensitivity after the saccades.

**Figure 1.**
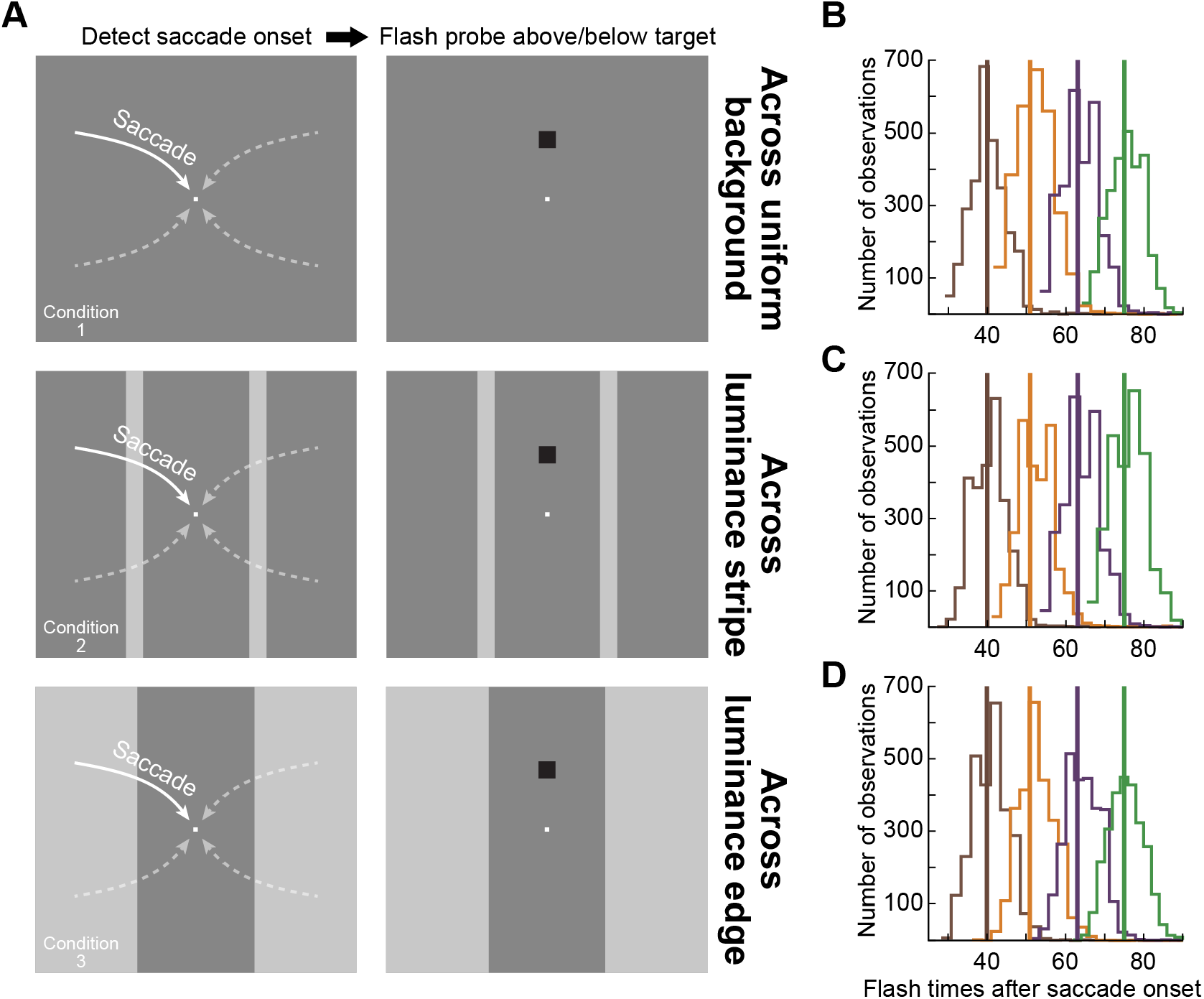
The different saccade conditions tested in our study. **(A)** Subjects made approximately 11.8 deg saccades towards display center from one of four possible locations. The saccades (schematized as curved arrows) were predominantly horizontal (see text), and we detected their onset online using a velocity criterion. Upon saccade detection, we presented a brief luminance flash at 7 deg either above or below the saccade target (right column of images). The flash was presented at one of four different delays from online saccade detection (see **B**-**D**). In condition 1 (top row), the saccades were made over a uniform background. In condition 2 (middle row), gaze crossed a vertical luminance stripe. And, in condition 3 (bottom row), the saccades brought gaze across a vertical luminance edge. We varied background luminance or the combination of pre- and post-saccadic background luminances across trials, and the probe flashes could also be of either positive (luminance increments relative to the background) or negative (luminance decrements) polarity. **(B)** After detecting saccades offline, we plotted the distributions of flash times collected during the experiments in condition 1 across all subjects. Our experimental manipulation succeeded in probing multiple times of perceptual sensitivity after saccade onset. The jitter in the individual distributions was due to variability in online saccade detection, as well as display update timing variability (due to the asynchrony between saccades and display refresh clocks). **(C, D)** Similar distributions of flash times for conditions 2 and 3, suggesting that differences in perceptual thresholds across conditions (see Results) were not attributable to different flash times relative to saccade onset. The thick vertical lines in **B**-**D** show the global average flash times for each colored distribution across all conditions.

Note that in all experimental conditions in which we used online saccade detection, we re-detected saccades offline in post-hoc analyses. This was because we could now refine saccade detection with an ability to look both forward and backward in time during offline analyses (Chen & Hafed, 2013); see the data analysis section below for further details. Across all experimental conditions with saccades being detected online, we confirmed that the 4 probe flash times designated above were occurring at 39.53 +/- 0.81, 51.38 +/- 0.85, 63.05 +/- 0.97, or 74.83 +/- 0.85 ms (mean value +/- s.e.m. across all trials from all subjects) after saccade onset (Fig. 1B-D shows the distributions of flash times in this and the two other saccade conditions described below). This was similar to the approach that we recently used (Idrees, Baumann, Franke, et al., 2020). Moreover, we confirmed that flash times after saccade onset were similar across all conditions employing real saccades (see raw distributions in Fig. 1B-D); this way, we could safely conclude that differences in the strength of perceptual saccadic suppression across the different conditions (see Results) could not be attributed to systematic differences in the flash times after saccades.

It should also be noted that with this approach of online saccade detection, we did not present flashes pre-saccadically. This was a conscious choice given the large numbers of trials needed in all of our experiments to collect full psychometric curves at each flash time and for each background luminance and for each of either saccade or fixation conditions. This is a strategy that we had also used in our recent experiments (Idrees, Baumann, Franke, et al., 2020). However, in other conditions described below (conditions 4 and 5), we did present flashes before visual transients, confirming that “pre-saccadic” suppression does occur (in an expected manner), and that it is consistent with all of our previously described results motivating the current study (J. Bellet et al., 2017; Chen & Hafed, 2017; Hafed & Krauzlis, 2010; Idrees, Baumann, Franke, et al., 2020).

The probe flash that was presented at different times after online saccade detection was a square of 2.4 by 2.4 deg dimensions (Idrees, Baumann, Franke, et al., 2020). In the current experiments, its luminance consisted of either a decrement (shown in the examples of Fig. 1A) or increment relative to the background luminance, allowing us to explore the impact of stimulus polarity on peri-saccadic perceptual sensitivity. The background luminance itself was either bright (77.3 cd/m^2^) or dark (22.4 cd/m^2^).

Across trials, the probe flash could either appear 7 deg above the saccade target location (i.e. above the intended saccade landing position) or 7 deg below the saccade target location, and the subjects’ task was to indicate, by pressing one of two buttons on a response box, whether the flash appeared above or below screen center in every trial. If the subjects did not see the flash, they had to guess its location. Note that with our display geometry, the retinotopic position at which the probe flash could appear was visually-stimulated before the saccade by the display itself (i.e. a uniform luminance) rather than the (very) dark surround of the laboratory. Thus, across the saccade, the possible retinotopic location of the flash was swept across a uniform luminance rather than across the outer edge of the display.

We collected psychometric curves of flash perception as follows. Across trials, and for a given condition (for example: the combination of dark background; negative contrast flash; flash time 24 ms after saccade detection), we used an adaptive approach to select flash luminance levels resulting in threshold performance (Idrees, Baumann, Franke, et al., 2020). Briefly, in the first of the two sessions for this condition (and similarly for all other conditions), we used a QUEST procedure (Watson & Pelli, 1983) for each flash polarity, each background luminance, and each post-saccadic flash time aiming to achieve a percent correct of 75% over 60 trials. Then, after the threshold contrast was found for each QUEST procedure, in the second session, we introduced, for each condition, 6 additional contrast levels around the detected perceptual threshold with the adaptive procedure, in order to obtain more samples for the psychometric curve of each subject. This is similar to what we did in our recent work (Idrees, Baumann, Franke, et al., 2020).

Note that in this and all other conditions and analyses throughout this study, we refer to “threshold” as the absolute value of the probe contrast that was used (whether the probe was a luminance increment or decrement relative to the background). However, we always also explicitly specify whether the probe was of positive (luminance increment) or negative (luminance decrement) polarity. This way, it is simpler to quantitatively compare saccade-related threshold elevations for both types of probe flashes.

We collected 1920 trials per subject in this condition.

#### (2) Real saccades across a vertical visual stripe

This condition was identical to condition 1 except that each saccade crossed a vertical luminance stripe of width 2.4 deg. The stripe was centered horizontally on the midway point between initial and final fixation spot locations, and it spanned the full vertical extent of the display (Fig. 1A, middle). Therefore, on every trial, the display was configured to have two vertical luminance stripes at 5.7 deg either to the right or left of display center. The left stripe was crossed by rightward saccades starting from the left side of the display, and the right stripe was crossed by leftward saccades. The rest of the display was the same as in condition 1. Across trials, we varied the luminance of either the background or the stripes. Specifically, the stripes had the dark luminance when the rest of the display had the bright luminance, and vice versa (same luminance values as in condition 1). This way, the eye could start and land on either a dark or bright luminance region (like in condition 1), but the difference is that the gaze would have had to cross a luminance stripe of the other luminosity during the saccade. Like in condition 1, note that with our display geometry, the retinal positions of the probe flash locations were visually-stimulated by the display before the saccade (rather than by the much darker surround in the laboratory environment around the display).

We collected 1920 trials per subject from this condition.

#### (3) Real saccades across a vertical luminance edge

In this condition, the display was split (along the horizontal dimension) into three areas (one central and two flanks; Fig. 1A, bottom): the central region had either a bright or dark luminance, and the flanks were the opposite. The vertical edge between the central region and either of the two flanking regions was at 5.7 deg horizontally from display center, which is halfway between the initial and final fixation spot’s horizontal locations. As a result, there was a vertical edge across which the same saccades as in conditions 1 and 2 were made. The bright and dark luminances on either side of the vertical edge were the same as those of the bright and dark background luminances in conditions 1 and 2, and they were varied across trials (some trials had the central area being bright, and others had the flanking areas being bright, independently of whether the saccade was rightward or leftward). Thus, in this condition, the saccade landed on either a bright or a dark luminance level (like in conditions 1 and 2); however, critically, the initial luminance level at saccade onset was always different from the final luminance level (Fig. 1A, bottom). Moreover, the probe flashes (whether luminance increments or luminance decrements) were always presented relative to the luminance level at the end of the saccade. That is, the probe flashes appeared on either bright or dark backgrounds (like in conditions 1 and 2).

We collected 1920 trials per subject from this condition.

#### (4) Simulated saccades across a vertical luminance edge

In this condition, the subjects maintained fixation at display center, and a visual transient like that in condition 3 was introduced. That is, the display had a central region that was either bright or dark and two flanking regions around it with the opposite brightness from the central region (Fig. 2A). Thus, there was a vertical edge, which was then translated horizontally to simulate a saccade across this edge. To achieve translation of the vertical edge across the fovea, we translated the whole displayed texture (i.e. both the central and flanking regions) horizontally at high speed (see the next paragraph for details on handling monitor edge effects). The resulting “simulated” saccade had parameters similar to those used in our recent work (Idrees, Baumann, Franke, et al., 2020). Briefly, we translated the texture (and, therefore, the vertical edge) by 1.9 deg every display frame (of approximately 12 ms) for 6 display frames, corresponding to an overall translation of 11.4 deg in 72 ms. This resulted in a high-speed translation of the vertical edge across the fovea during maintained fixation. The translation matched, in amplitude and duration, the amplitudes and durations of the real saccades that the subjects made in conditions 1-3 above. Specifically, in these conditions, the average saccade amplitude and duration were 11.37 +/- 0.14 deg and 74.89 +/- 2.05 ms, respectively, across all trials from all subjects (mean value +/- s.e.m.).

**Figure 2.**
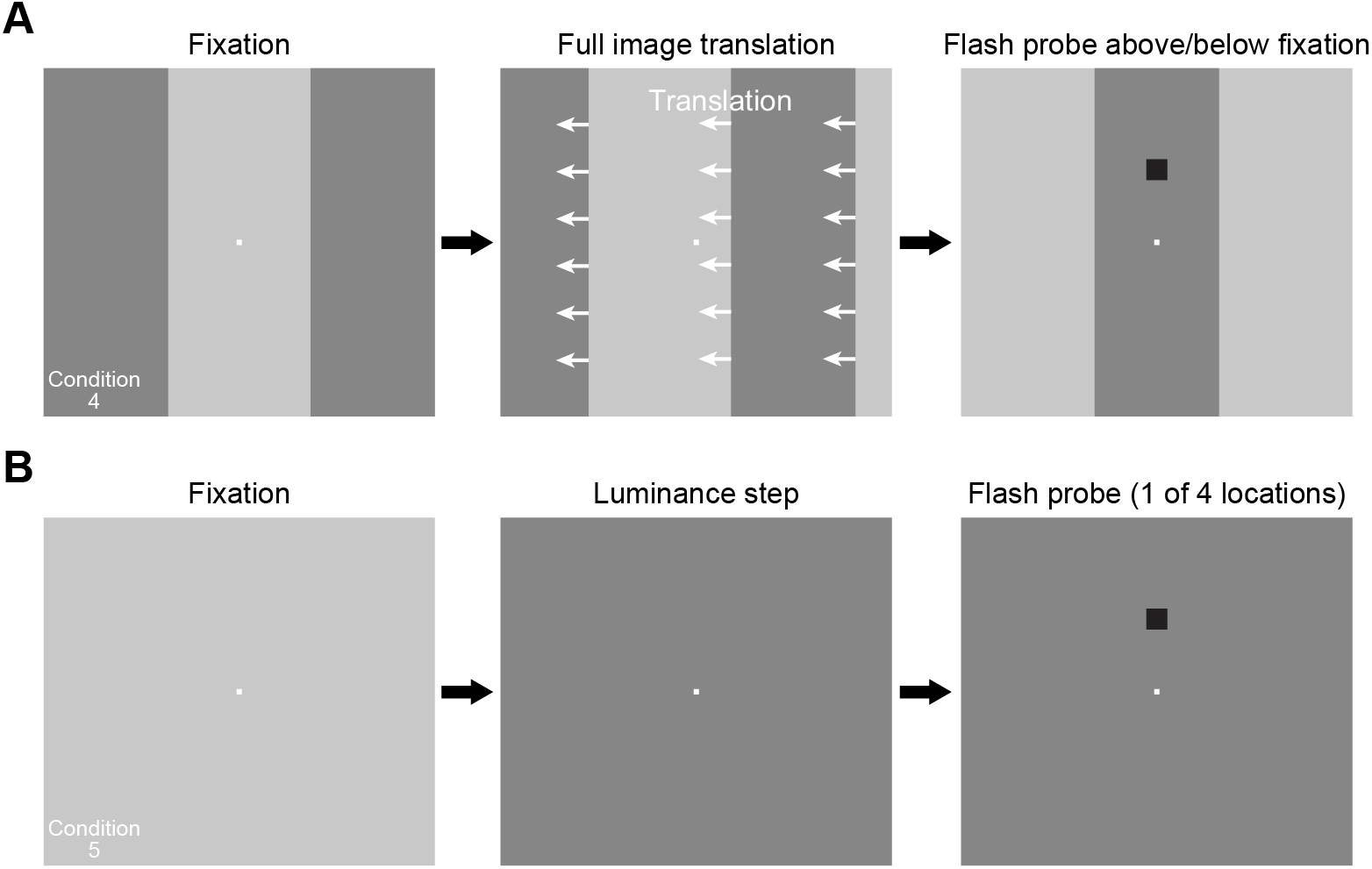
The visual-only conditions tested in our study. **(A)** In condition 4, subjects fixated the center of the display, and the background was identical to that of condition 3 in Fig. 1. After initial fixation was established, the whole background translated horizontally to shift one of the two vertical edges across gaze position. After the translation, gaze was now aimed at a different background luminance. This condition mimicked the real saccade version in condition 3. Note that during translation, we treated the texture as being horizontally periodic (in the shown example, this meant that a bright background was revealed on the right edge of the display monitor during leftward translation). At different times relative to translation onset, we presented a flash either above or below fixation, similar to Fig. 1A. In this case, the flashes could be presented either before or after translation onset (Methods). When presented after translation onset, the timing was such that they were presented over the new background luminance after translation end (that is, the edge had crossed display center even for the earliest flash time after translation onset). **(B)** In condition 5, the subjects fixated the center of a uniform display, and we changed the luminance of the background (from dark to bright or vice versa). Relative to this transient, a probe flash could happen at similar times to those used in **A**. Note that in this condition only, we had four possible probe flash locations, because this experiment was a replication of our earlier version of it in (Idrees, Baumann, Franke, et al., 2020; Idrees, Baumann, Korympidou, et al., 2020).

The sequence of events in a given trial was as follows. An initial fixation spot appeared at trial onset at the center of the display. The two vertical edges were at 5.7 deg to the left and right of the fixation spot location (Fig. 2A). After a random time of 800-1700 ms, the high-speed translation was started. For a leftward translation, the right vertical edge shifted to become now located at the initial location of the left vertical edge after the entire sequence had ended, and vice versa for a rightward texture shift. To handle display monitor edge effects, we treated the initial displayed pattern (i.e. the whole of the combination of central region and two flanks) as a horizontally periodic pattern. As a result, when a leftward shift happened, the rightward flank translated to become the central region, and a new rightward flank that was the same luminance as the original central region appeared. The corresponding scenario played out for a rightward shift. A probe flash appeared similar to the other conditions described above.

Across trials, we picked 2 times of probe flashes after the onset of the simulated saccades (47 and 59 ms) and also 2 times of probe flashes before the onset of the simulated saccades (24 and 35 ms before simulated saccade onset). The use of probe flashes before simulated saccade onset allowed us to demonstrate existence of pre-transient suppression, which provides a logical link to our other work (Idrees, Baumann, Franke, et al., 2020), and which also demonstrates that our lack of pre-saccadic flash times in conditions 1 to 3 above does not necessarily mean that there was no pre-saccadic suppression in these conditions. All other parameters of the experiment were similar to conditions 1 to 3.

We collected 1920 trials per subject from this condition.

#### (5) Fixation with a luminance change

In this final condition, subjects maintained fixation at the center of a uniform display, and the visual transient now consisted of a change in background luminance from dark to bright, or vice versa (Fig. 2B). Probe flashes could appear either before (24 or 35 m) or after (35, 71, or 106 ms) the luminance change, like in condition 4. Note that in this condition only, we used a four-alternative forced choice paradigm as opposed to the two-alternative forced choice paradigm in all other conditions. So, instead of just presenting a probe flash 7 deg above or below the fixation (or saccade) target location, the probe flash could now appear at 7 deg in any of the four cardinal directions around the fixation spot (right, left, up, or down; Fig. 2B). Subjects had to press one of four buttons corresponding to the four possible flash locations. The reason that we used four alternatives in this case is that fixation was always at the center of the display and that the display itself was uniform (allowing us to probe horizontal flash locations without worrying about “pre-saccadic” retinal image regions of horizontal flash locations being visually stimulated by the dark laboratory surround instead of the actual display, or by another luminance on the display like in condition 4). Another reason for using four alternatives is that this condition involved collecting new data from a similar condition (with four alternatives) that we had recently run (Idrees, Baumann, Franke, et al., 2020; Idrees, Baumann, Korympidou, et al., 2020). All other parameters and procedures were the same as in all other conditions.

We collected 2400 trials per subject from this condition.

#### Data analysis

We detected saccades and microsaccades using our laboratory’s established methods (M. E. Bellet, Bellet, Nienborg, Hafed, & Berens, 2019; Chen & Hafed, 2013; Hafed, 2013). For all trials with fixation (conditions 4 and 5), we ensured that there were no microsaccades from -200 to +50 ms relative to probe flash onset. This allowed us to avoid potential changes in visual sensitivity around the time of microsaccades (Chen & Hafed, 2017; Chen et al., 2015; Hafed, Chen, & Tian, 2015). Trials that contained microsaccades in the above intervals were excluded from subsequent analyses. Similarly, for trials with real saccades, we excluded trials with saccades before instruction (i.e. before fixation spot jump) and also trials with saccadic reaction times >500 ms from saccade target onset, and we confirmed that the flash times relative to saccade onset (after post-hoc offline saccade detection) indeed clustered into 4 different time points, as designed in the task (see above and Fig. 1B-D).

We analyzed the proportion of correct trials as a function of probe flash Weber contrast (i.e. absolute value of luminance difference of flash from background luminance, divided by the background luminance). We did this for each flash luminance used, and independently for whether the probe flash consisted of a luminance increment or a luminance decrement relative to background luminance. We also did this for each flash time used (for real saccades, we classified all flash times into 4 “clusters” of flash times centered around the mean values measured after offline saccade detection; see Fig. 1B-D). Across contrasts (for either increments or decrements), we obtained a psychometric curve fit of perceptual performance using the psignifit 4 toolbox (Schutt, Harmeling, Macke, & Wichmann, 2016). Briefly, we used, via the toolbox, a beta-binomial model for the psychometric curve. We defined the perceptual threshold as the absolute Weber contrast value resulting in a correct performance level that was at either 75% of the total dynamic range of the fitted psychometric curve (for conditions 1-4 with two-alternative forced choices) or 62.5% of the total dynamic range of the psychometric curve (for condition 5 with four-alternative forced choices). If there was perceptual suppression of sensitivity, then the threshold contrast was elevated. Thus, plots of threshold contrast indicate maximal suppression when the perceptual thresholds are high.

For each background luminance over which a given flash appeared (dark or bright), and for each flash time, we calculated a psychometric curve for either flashes consisting of luminance increments or luminance decrements (i.e. flash stimulus polarity). We used the longer flash times after visual transient onset (either caused by real saccades or visual display updates during fixation) to confirm that there was perceptual recovery with time (as expected). Therefore, elevations of threshold contrast above the threshold values at such longer post-transient times were evidence for perceptual suppression.

To summarize results across subjects, we first estimated perceptual thresholds for each subject, and we then averaged the thresholds across subjects, with indications of inter-subject variance in all figures. To perform statistics, we used two-way ANOVA’s testing the influences of flash time and condition on perceptual thresholds. We also sometimes performed individual t-tests comparing pairs of conditions at one given flash time; typically, this was the shortest flash time after saccade onset (or after texture translation or contrast change in conditions 4 and 5) since this time was the time associated with maximal perceptual suppression. This flash time was therefore of most interest when comparing suppression strength across the different conditions. It should also be noted that in conditions 4 and 5, when we had “pre-transient” flashes, the flash appeared on a background that was different from the background for flash onsets occurring after the visual transient. Therefore, in all figures, we always pooled data points based on the background luminance on which a flash actually appeared. For example, pre-transient flashes in condition 4 with, say, a translation from bright to dark background were plotted as flashes over a bright background, whereas post-transient flashes were plotted as flashes over a dark background. This way, we always compared perception with a similar relationship between flash luminance and background luminance at the time of the flash. In statistical analyses of condition 5, we also pooled, within each subject, the longest two flash times after background luminance change together. This was because these two times showed the same effects anyway.

## Results

We assessed the visual contributions to perceptual saccadic suppression by measuring perceptual sensitivity for brief flashes presented around the time of saccades across a variety of image types. We found that perceptual saccadic suppression has a large component of visual dependencies built into it. Critically, we also compared such sensitivity to situations in which there were no saccades, but in which conceptually comparable visual transients like those associated with saccades were experimentally introduced. With such simulated saccades, we found similar suppression as during real saccades, despite the absence of saccadic motor commands, and the same dependencies on visual context. In all cases, we compared perceptual sensitivity to a baseline case in which saccades were made across a uniform background. These results confirm and extend our recent observations that were motivated by recordings of ex-vivo retinal ganglion cell activity in multiple species (Idrees, Baumann, Franke, et al., 2020; Idrees, Baumann, Korympidou, et al., 2020), and they are in line with other psychophysical evidence in the literature. This allows us to conclude that perceptual saccadic suppression may fundamentally be a visual phenomenon. In what follows, we describe how we reached this conclusion.

### Saccades across a local luminance feature are associated with much stronger perceptual suppression than saccades across a uniform background

We collected perceptual threshold measurements across saccades. In one condition (condition 1), subjects made predominantly horizontal oblique saccades across a uniform background. In another condition, the saccades were made across a 2.4 deg wide luminance stripe relative to background luminance, but the luminances at both saccade start and saccade end were the same (condition 2). That is, the display consisted of a uniform background with a vertical stripe (of 2.4 deg width) midway between the start and end points of the saccade (see Fig. 1). We detected saccade onset online, and we presented probe flashes at 4 different times after saccade detection (Methods). The presented peri-saccadic probe flashes occurred on top of the same background luminance in both the baseline control condition (saccades across a uniform background) as well as the luminance feature condition (saccades across a luminance stripe on an otherwise uniform background). Subjects had to decide, on every trial, whether the probe flash appeared above or below the line of site (Methods), and we varied flash contrasts in order to collect full psychometric curves.

Figure 3 shows example psychometric curves of perceptual detectability obtained from an example subject in the two conditions and at flash times of approximately 40 ms (Fig. 3A) and 63 ms (Fig. 3B) after saccade onset. In this figure, we show results only from the dark background condition (Methods), and also when the luminance of the probe flash was a decrement relative to background luminance (i.e. negative stimulus polarity). Note that in this and all other figures, we depict the absolute value of Weber contrast, to aid comparison of effects between positive-contrast and negative-contrast probe flashes. Flashes immediately after saccade detection (Fig. 3A) were associated with elevated perceptual thresholds in both conditions: this is evidenced by rightward shifts of the psychometric curves for the early time compared to the late one (compare the correspondingly colored curves in Fig. 3A and Fig. 3B). However, the effect was much stronger for condition 2, in which saccades crossed over a bright luminance stripe of 2.4 deg width. Specifically, at 40 ms (Fig. 3A), the absolute value of perceptual threshold was at >0.8 Weber contrast in condition 2 and at only <0.15 Weber contrast in condition 1. Similarly, at the later time point, even after partial perceptual recovery (Fig. 3B), the absolute value of threshold in condition 2 was still at a level of almost 0.3 Weber contrast, but it was already at a level <0.09 in condition 1. Thus, perceptual saccadic suppression is much stronger when saccades are made across a luminance stripe than across a uniform background.

**Figure 3.**
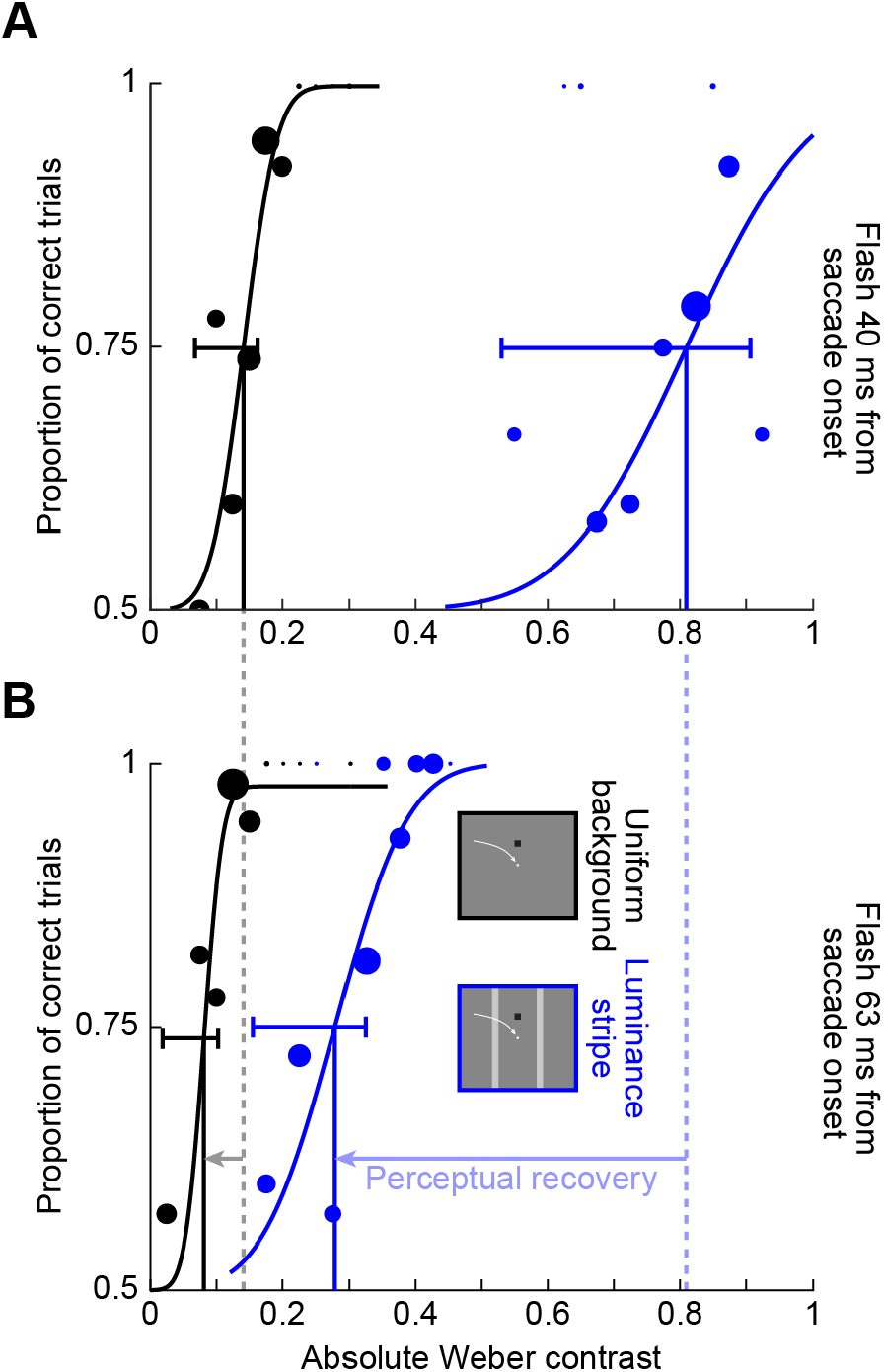
Much stronger saccadic suppression across a luminance stripe than across a uniform background. Example psychometric curves of perceptual detectability from one sample subject, for dark probe flashes over a dark background. Circles show individual data points, with the size of the circle scaled based on the numbers of observations collected. The blue curves show psychometric curves when the saccade crossed a luminance stripe (condition 2), whereas the black curves show the results obtained when the saccade was made across a uniform background (condition 1); see the schematic legend in **B**. Vertical lines indicate the perceptual threshold values in each condition, and horizontal error bars show 95% confidence intervals for each psychometric curve (obtained using the psignifit 4 toolbox; Methods). **(A)** Data for probe flashes presented immediately after online saccade detection (40 ms after actual saccade onset; first time cluster in Fig. 1B-D). There was a much larger rightward shift in the psychometric curve in the case of condition 2 when compared to condition 1. **(B)** Data for a later flash time (63 ms after saccade onset; third time cluster in Fig. 1B-D). Both curves recovered at this time point (i.e. shifted leftward relative to **A**; horizontal leftward arrows), but the blue curve still showed much more suppression (i.e. rightward shift) compared to the black curve.

Across subjects, we made very similar observations. For example, the blue curve in Fig. 4A shows the average psychometric curve obtained across all subjects (the average of all subjects’ individual psychometric curves) in condition 2 for the first flash time (40 ms after saccade onset). This curve was shifted much more strongly to the right than the black curve in the same panel. The black curve summarizes the results across subjects with only a uniform background (condition 1). Similarly, Fig. 4B shows recovery results at the 63 ms time point in the same format, again consistent with Fig. 3. To summarize the effect size and time course to recovery across the two conditions, we then estimated the perceptual threshold for a given condition as the absolute value of Weber contrast in each psychometric curve resulting in a correct performance level of 75% of the dynamic range (Methods, compare the vertical lines in Fig. 3) (Idrees, Baumann, Franke, et al., 2020). Figure 4C plots the absolute threshold contrasts at all probe flash times: in blue for condition 2 and in black for condition 1. Error bars denote s.e.m. across subjects (Methods). As can be seen, there was much stronger suppression (larger increase in the perceptual threshold contrast) for saccades across the luminance stripe (blue) when compared to saccades across the uniform background (black), and this also persisted as a function of time. Statistically, a two-way ANOVA revealed a highly significant effect of both factors of flash time (F(3,40) = 43.3, p <10^−11^) and condition (F(1,40) = 151.1, p <10^−14^). Moreover, there was also a significant interaction between the two factors (F(3,40) = 29.2, p < 10^−9^). Post hoc, we compared the thresholds of condition 1 (mean +/- s.e.m.: 0.13 +/- 0.02 Weber contrast) and condition 2 (0.95 +/- 0.1 Weber contrast) at 40 ms (the time of maximal suppression in our experiments). There was a highly significant difference around the time of maximal suppression (t(5) = 7.9, p < 0.001).

**Figure 4.**
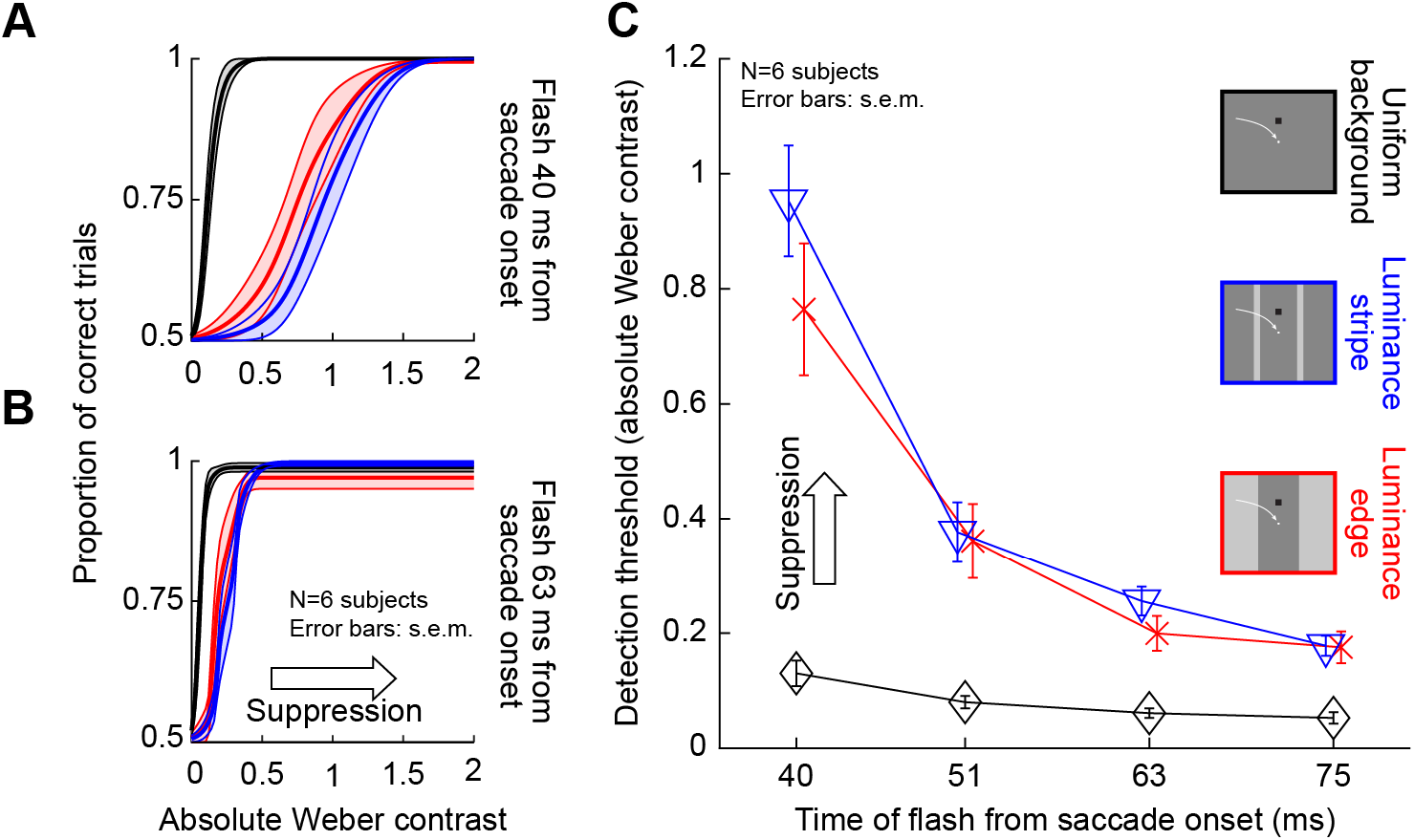
Systematically stronger saccadic suppression across a luminance stripe or luminance edge than across a uniform background. **(A)** Average psychometric curves of perceptual detectability from all subjects when the probe flashes occurred 40 ms after saccade onset (first time cluster in Fig. 1B-D). Black: psychometric curve for condition 1, when saccades were made across a uniform background. Blue: psychometric curve when saccades crossed a luminance stripe (condition 2). Red: psychometric curve for condition 3, when saccades crossed a luminance edge. In all cases, error bands denote s.e.m. across subjects. Perceptual thresholds were elevated much more for conditions 2 and 3 than for condition 1. **(B)** Same as in **A** but for a later time point (63 ms). There was perceptual recovery, but saccadic suppression was still stronger in conditions 2 and 3 than in condition 1; the subjects always had higher thresholds than in condition 1 (also see **C**). **(C)** Perceptual thresholds (average across subjects) for each flash time after saccade onset. Higher values mean stronger saccadic suppression. Error bars denote s.e.m. across subjects. Note that the time values on the x-axis are the average flash times obtained after offline saccade detection (Methods and Fig. 1B-D). Also note that for each average flash time, we slightly jittered the x-axis position of each data point to not mask individual data points and their error bars (also true in subsequent figures below). There was much stronger saccadic suppression in conditions 2 and 3 than in condition 1.

### Different pre- and post-saccadic visual stimulation is also associated with strong perceptual saccadic suppression

We next asked whether the final luminance on which the probe flashes appeared peri-saccadically needed to be the same as the “pre-saccadic” luminance at the line of sight for us to obtain the above results. To test this, we replaced the stripe used in condition 2 with a luminance edge (condition 3; Methods; red curves in Fig. 4). That is, this time, the subjects initially fixated a bright background, and they then made saccades across a vertical edge, such that the probe flashes (and saccade landing positions) were now over a dark background. We then compared perceptual suppression to that obtained with saccades across a uniform dark background (condition 1). The same strong difference in saccadic suppression relative to the uniform dark background was observed (Fig. 4, now compare red to black curves). For example, at the first flash time after saccade onset (Fig. 4A, C), the average perceptual threshold in both conditions 2 and 3 was >0.75 Weber contrast, whereas it was only 0.13 Weber contrast when saccades were made across a uniform background (Fig. 4C). Moreover, statistically, there was still a highly significant effect of time (F(3,40) = 19, p <10^−7^) and condition (F(1,40) = 70, p <10^−9^), when now comparing condition 3 to condition 1 (two-way ANOVA) instead of comparing condition 2 to condition 1 as we had done above. There was also a significant interaction effect (F(3,40) = 11.4, p <10^−5^). Moreover, a paired post hoc t-test highlighted a difference at at 40 ms (t(5) = 6.72, p = 0.001); condition 1: 0.13 +/- 0.02 s.e.m. Weber contrast, and condition 3: 0.76 +/- 0.11 s.e.m. Weber contrast. In fact, when we compared threshold contrasts across subjects in both conditions 2 and 3 together (colored curves in Fig. 4C), we found that suppression was equally strong for both of these conditions. A two-way ANOVA with the factors time and condition (2 versus 3) revealed no significant interaction between the factors (F(3,40) = 0.9, p = 0.45) and no significant effect of the condition (F(1,40) = 2.2, p = 0.15). However, expectedly, the effect of flash time was highly significant (F(3,40) = 48.5, p <10^−12^).

Therefore, saccadic suppression is also enhanced when gaze crosses a luminance edge rather than a luminance stripe, replicating results from an earlier study (Maij, Matziridi, Smeets, & Brenner, 2012). The difference here is that in our current experiments, we assessed thresholds explicitly, by collecting full psychometric curves, allowing a more sensitive measure of perceptual sensitivity. We also varied stimulus polarities of the flashes, and even replaced saccades with image translations, as we describe in more detail shortly.

It is important to note that saccadic suppression still occurred over a uniform background (condition 1), albeit in a weak fashion. To confirm this, we statistically assessed perceptual detectability in condition 1 with a dark background, and we found that there was elevation of contrast immediately after saccade detection when compared to the latest two flash times after saccade onset in Fig. 4C (black): a one-way ANOVA with flash time as factor was significant (F(3,20) = 6.2, p = 0.004), and a post hoc test revealed significant differences for the perceptual thresholds at 40 and 63 ms (p = 0.01) and 40 and 75 ms (p =0.004). Similar results also occurred for a bright background (we address the influence of background luminance more explicitly shortly). This extends the results of (Maij et al., 2012) in an important way. Specifically, in that study, it could appear that there was no saccadic suppression at all with a uniform background. However, we think that measuring contrast thresholds in our experiments was a more sensitive measure of perceptual sensitivity. The approach of Maij et al could result in floor or ceiling effects, potentially masking a mild amount of suppression (which we observed).

### A visual dependence of saccadic suppression also emerges when considering the luminance of the background across which saccades are made

The above results indicate that the visual conditions across which saccades are made do matter for perceptual saccadic suppression, as we also recently argued (Idrees, Baumann, Franke, et al., 2020). To further demonstrate that perceptual saccadic suppression does indeed have a strong visual component, we next checked whether the luminance of the background mattered for the strength of suppression. In our experiments, saccades could be made in condition 1 across either a bright or a dark background (Methods). Similarly, in conditions 2 and 3, the final landing position of the saccades, and therefore the final possible probe flash locations, could be either on a bright background or a dark background. In the description of results so far we have focused only on the dark background condition. We now describe the threshold contrasts that were obtained when the probe flashes appeared on a bright background instead.

Fig. 5A, B show average psychometric curves across subjects for the first flash time after saccade detection (approximately 40 ms from saccade onset). Here, we directly compare trials having a dark background at flash occurrence (solid lines, replicated from Fig. 4A) with trials having a bright background at flash occurrence (dashed lines). In Fig. 5A, the data for condition 2 are shown, and in Fig. 5B, the data for condition 3 are shown. In both cases, saccadic suppression was much stronger over a dark landing position than over a bright landing position. This effect can be better seen through plots of the time course of saccadic suppression in the two conditions (Fig. 5C). Statistically comparing bright versus dark backgrounds, as well as time, in two-way ANOVA’s confirmed that there was a strong effect of background luminance (and time) on saccadic suppression. For both conditions, a two-way ANOVA revealed significant interactions (condition 2: F(3,40) = 20.9, p <10^−7^; condition 3: F(3,40) = 7.1, p <0.001) between the factors, as well as highly significant main effects for time (condition 2: F(3,40) = 52.9, p < 10^−13^; condition 3: F(3,40) = 24.4, p <10^−8^) and background luminance (condition 2: F(1,40) = 102.2, p <10^−11^; condition 3: F(1,40) = 24.2, p <10^−4^). Note that for simplicity, we still only considered flashes consisting of luminance decrements relative to the background (i.e. of negative stimulus polarity), exactly like in all analyses above. We will explicitly describe the impact of flash luminance polarity relative to the background in a subsequent analysis. In any case, threshold elevations were much stronger with dark backgrounds than with bright backgrounds.

**Figure 5.**
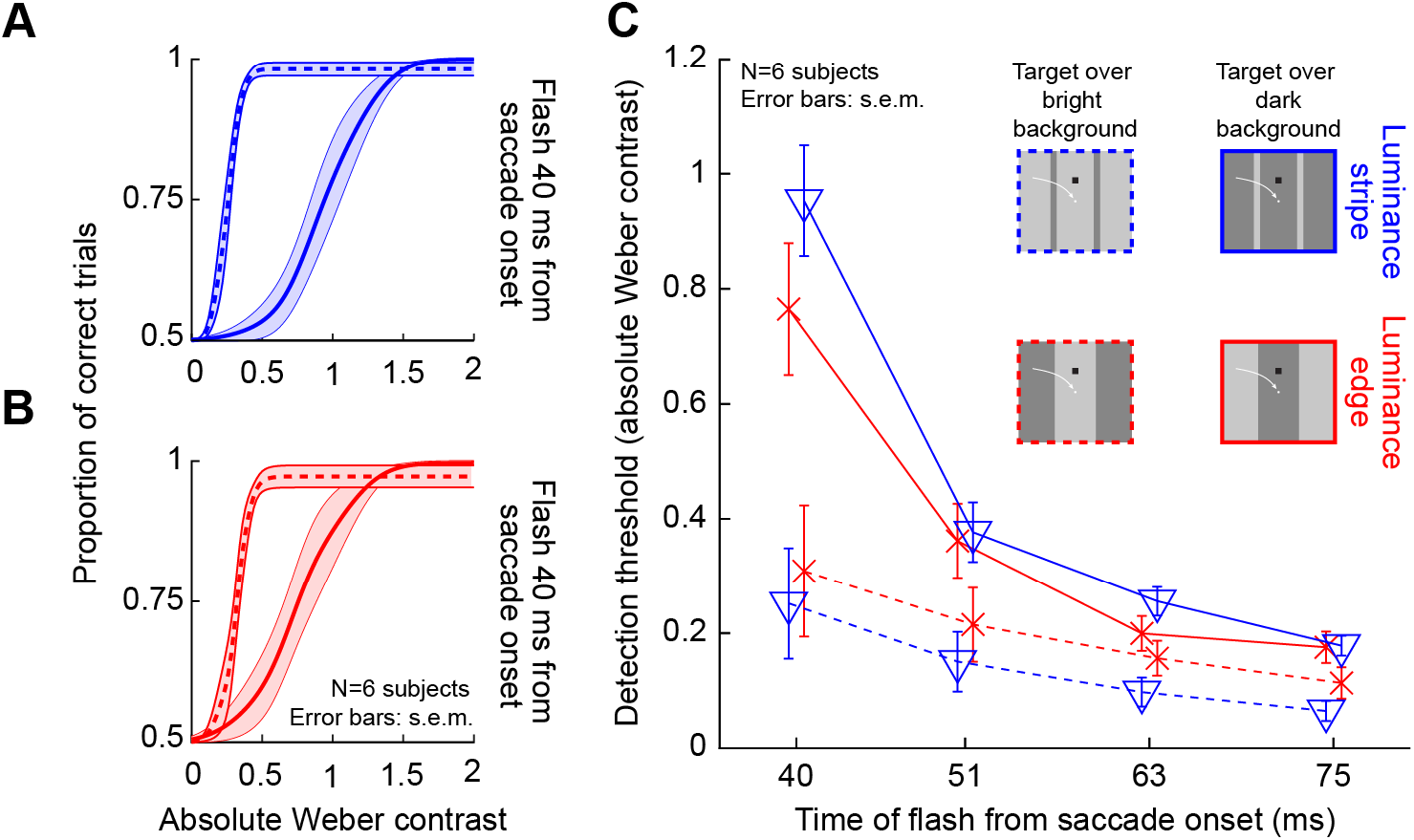
Stronger saccadic suppression over dark backgrounds, whether with luminance edges or luminance stripes. **(A)** Average psychometric curves of perceptual detectability from all subjects in condition 2, and for probes occurring 40 ms after saccade onset (i.e. the first time cluster in Fig. 1B-D). Solid indicates flashes over a dark background (i.e. saccades starting and ending on a dark patch, but crossing a bright stripe), and dashed indicates flashes over a bright background (i.e. saccades starting and ending on a bright patch, but crossing a dark stripe). In both cases, the probe flashes had negative luminance polarity (i.e. they were luminance decrements relative to the background). Saccadic suppression was stronger with dark backgrounds than with bright backgrounds. Error bands denote s.e.m. across subjects. **(B)** Similar results were obtained in condition 3. Flashes over a dark background (when saccades were made from a bright to a dark patch) were associated with stronger suppression than flashes over a bright background (when saccades were made from a dark to a bright patch). **(C)** Perceptual thresholds (averaged across subjects) for each flash time after saccade onset. Higher values mean stronger saccadic suppression, and error bars denote s.e.m. across the subjects. In both conditions 2 and 3, flashes over a dark background (solid) were associated with much stronger saccadic suppression than flashes over a bright background (dashed). All other conventions are similar to Fig. 4.

Incidentally, for saccades across a uniform background (condition 1), we did not observe a significant influence of background luminance on the strength of saccadic suppression. A two-way ANOVA with the factors time and background luminance showed no interaction between the factors (F(3,40) = 0.05, p = 0.98). As expected, since suppression still took place in this condition, the main effect of time was still significant (F(3,40) = 13.5, p < 10^−5^). However, the factor of background luminance was not significant (F(1,40) =0.9, p = 0.34). This could reflect the small effect size in this condition in general.

Therefore, the results of Fig. 5 further illustrate the strong visual dependencies of perceptual saccadic suppression. When (negative polarity) probe flashes were presented over dark backgrounds, perceptual thresholds were elevated much more around the time of saccades than when the same negative polarity flashes (of the same contrast) were presented over bright backgrounds. Earlier work related to condition 3 did not analyze the effects of background luminance (Maij et al., 2012), and our recent experiments only used textured backgrounds (Idrees, Baumann, Franke, et al., 2020). Interestingly, the present results clarify why saccadic suppression was so strong in these recent experiments (Idrees, Baumann, Franke, et al., 2020). This was because a visual feature seems to be necessary for boosting the saccadic suppression effect.

### Flash luminance polarity interacts with background luminance to modulate the strength of saccadic suppression

We observed above that saccades across luminance-defined features are associated with strong perceptual suppression (Figs. 3, 4), and that background luminance interacts with the presented brief flashes to modulate the strength of suppression (Fig. 5). However, in all of our analyses so far, we only considered negative polarity probe flashes (that is, flashes consisting of luminance decrements relative to the surrounding background luminance). To check whether probe flash polarity constitutes an additional visual dependence of perceptual saccadic suppression, we then turned our attention to analyzing trials with positive polarity probe flashes (that is, flashes that had luminance increments relative to the surrounding background luminance).

In Fig. 6, we plotted the threshold contrasts across subjects for positive and negative polarity flashes in conditions 1, 2, and 3. In Fig. 6A, C, we show the results with flashes presented over a dark background, and in Fig. 6B, D, we show the results with flashes presented over a bright background. The top row of the figure (Fig. 6A, B) shows results from condition 2 (blue), whereas the bottom row (Fig. 6C, D) shows results from condition 3 (red). For each condition, the different shades of color denote whether the flash had negative polarity (saturated colors; these are data replicated from the previous figures) or positive polarity (unsaturated colors) relative to the background. Note that both rows also show the results from condition 1 with a uniform background for reference (black and gray curves). Also note that the y-axes in Fig. 6A, C are different from those in Fig. 6B, D because of the differential impact of background luminance that we described in Fig. 5. As can be seen from Fig. 6A, C, negative polarity probe flashes over dark backgrounds had stronger saccadic suppression (higher detection thresholds) than positive polarity probe flashes over the same background, especially at the time of peak saccadic suppression (40 ms). For example, immediately after saccade onset in conditions 2 and 3, perceptual thresholds were at 0.95 and 0.76 Weber contrast across subjects for negative polarity flashes, respectively (Fig. 6A, C, colored curves). For positive polarity flashes, the thresholds were elevated to only 0.69 and 0.54, respectively. Statistically, this difference between positive and negative polarity flashes was significant at 40 ms in both condition 2 (t(5) = 6.69, p = 0.001; positive polarity mean and s.e.m.: 0.95 +/- 0.1 Weber contrast; negative polarity: 0.69 +/- 0.07 Weber contrast) and condition 3 (paired t-test: t(5) = 3.31, p = 0.02; positive polarity mean and s.e.m.: 0.76 +/- 0.11 Weber contrast; negative polarity: 0.54 +/- 0.07 Weber contrast). Interestingly, over a bright background (Fig. 6B, D), both conditions 2 and 3 did not show any difference in the strength of saccadic suppression between positive and negative polarity flashes (Fig. 6B, D, colored curves). Therefore, there was an interaction between flash polarity and background luminance in modulating the strength of perceptual saccadic suppression.

**Figure 6.**
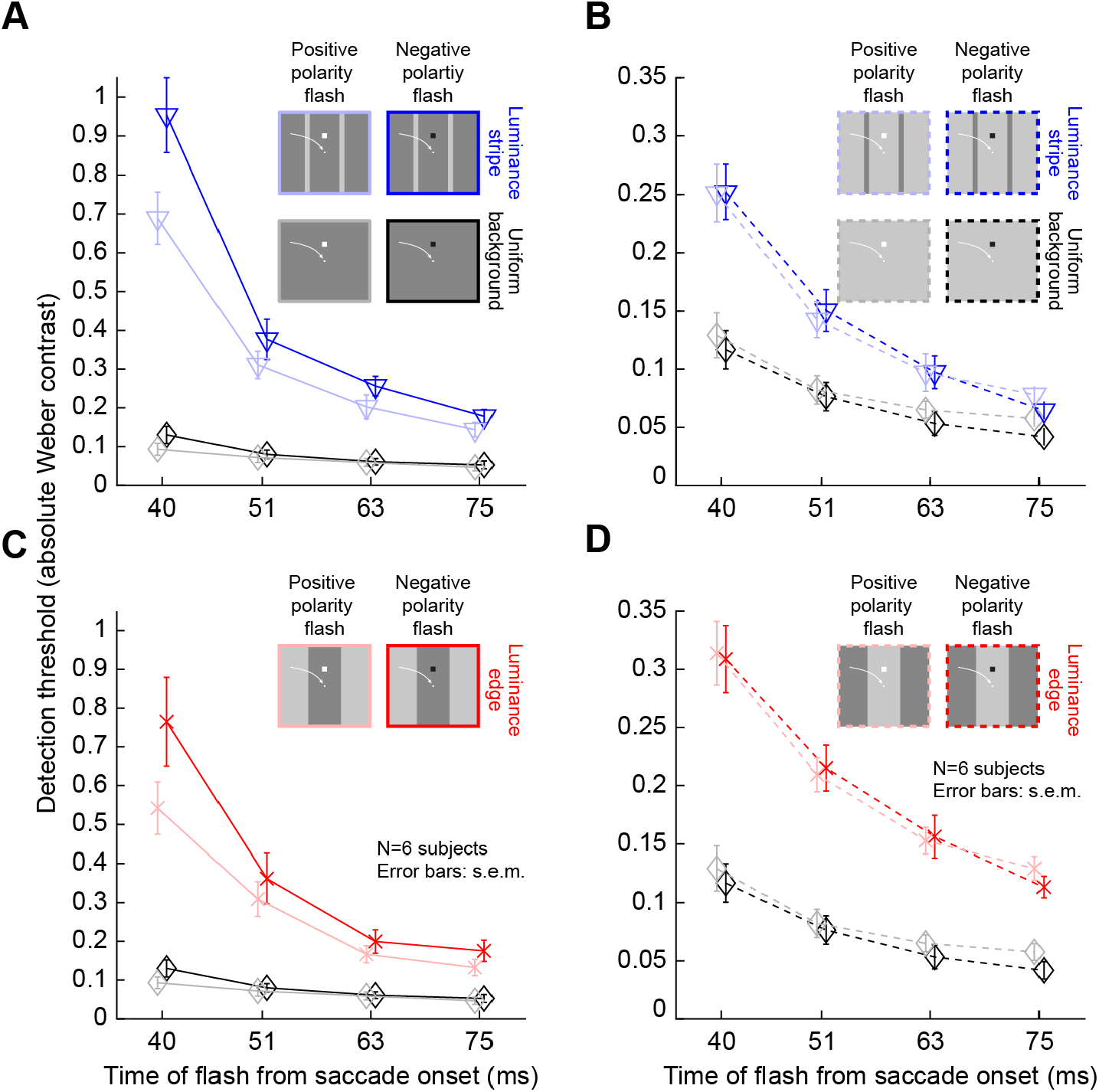
Stronger saccadic suppression for negative polarity flashes over dark backgrounds when compared to positive polarity flashes. Perceptual thresholds across subjects as a function of flash time after saccade onset (error bars denote s.e.m. across subjects). **(A)** Data from condition 2 (blue) and condition 1 (black) with flashes appearing over a dark background. The saturated colors show results with negative polarity flashes, and the unsaturated colors show results with positive polarity flashes. Consistent with earlier figures, perceptual saccadic suppression was much stronger (higher absolute values of perceptual thresholds) in condition 2 than in condition 1 (compare colored to black curves). Within each condition, saccadic suppression was stronger with negative polarity flashes than positive polarity flashes, especially near the time of peak saccadic suppression (40 ms). **(B)** With bright backgrounds, the effects of flash polarity were absent (but suppression was still stronger than in condition 1). Note that the y-axis is different from **A**, because suppression was weaker for flashes over bright backgrounds (see Fig. 5). **(C, D)** Similar results from condition 3, with saccades crossing a luminance edge. The black curves in this case are the same as in **A, B**. All other conventions are similar to Figs. 4, 5.

It is also interesting that for condition 1, similar effects still existed for flash luminance polarity (Fig. 6, black and gray curves) despite the very weak perceptual saccadic suppression that occurred: it was still the case that negative polarity flashes over a dark background (mean threshold 0.13 +/- 0.02 s.e.m. Weber contrast) tended to cause stronger saccadic suppression than positive polarity flashes (mean threshold: 0.09 +/- 0.01 s.e.m.) over the same background (t(5) = 3.6, p = 0.016 Weber contrast). Thus, our analyses of probe flash stimulus polarity in all saccade conditions (1, 2, and 3) provide further evidence that perceptual saccadic suppression has strong visual dependencies.

### Perceptual suppression occurs equally well, and with the same visual dependencies, for image sweeps that conceptually mimic saccade-induced image shifts

The above evidence is indicative of strong and rich visual-visual interactions in saccadic suppression between the saccade-related retinal image shifts and the probe flashes themselves, as we have recently suggested (Idrees, Baumann, Franke, et al., 2020; Idrees, Baumann, Korympidou, et al., 2020). If that is indeed the case, then the same visual dependencies as those demonstrated in Figs. 3-6 should also happen in the complete absence of saccades, but in the presence of saccade-like image shifts and probe flashes. We therefore repeated the above experiments but with so-called “simulated saccades”. We asked subjects to fixate, and we swept a vertical luminance-defined edge across the retina to simulate a saccade-like image displacement (Fig. 2A). This condition was therefore conceptually similar to condition 3 (Methods). The probe flashes happened identically to how they happened in the real saccade version of the experiment; that is, they occurred over a dark or bright background, and they also had either positive (luminance increment) or negative (luminance decrement) stimulus polarity. We also additionally tested pre-shift probe flashes, to check for perceptual suppression before “simulated saccade” onset (Idrees, Baumann, Franke, et al., 2020).

We replicated all of the above observations that were made with real saccades. First, we confirmed that immediately after saccade-like image translation, strong perceptual suppression occurred, which was much stronger than the suppression with real saccades across a uniform background. This effect is demonstrated in Fig. 7A, which shows the average psychometric curve across all subjects when a negative polarity flash appeared over a dark background at 47 ms after image translation onset (magenta curve). For reference, the black curve in the same figure shows the average psychometric curve associated with “real” saccadic suppression over a uniform (dark) background at our shortest flash time (i.e. associated with maximal saccadic suppression in our data set). As can be seen, perceptual thresholds were strongly elevated even with the simulated saccades, just like in condition 3. Across flash times, Fig. 7B shows that immediately after image shift onset, the absolute value of threshold contrast over a dark background was 0.54 Weber contrast for negative polarity flashes. This value was much higher than that for real saccades across uniform backgrounds (0.13 Weber contrast for the shortest flash time with the real saccade; 40 ms). Statistically, we confirmed this by doing a paired comparison of thresholds between condition 4 (0.54 +/- 0.11 s.e.m. Weber contrast) and condition 1 (0.13 +/- 0.02 s.e.m. Weber contrast) at the shortest positive flash time after either image translation onset or saccade onset (t(5) = 4.7, p = 0.005).

**Figure 7.**
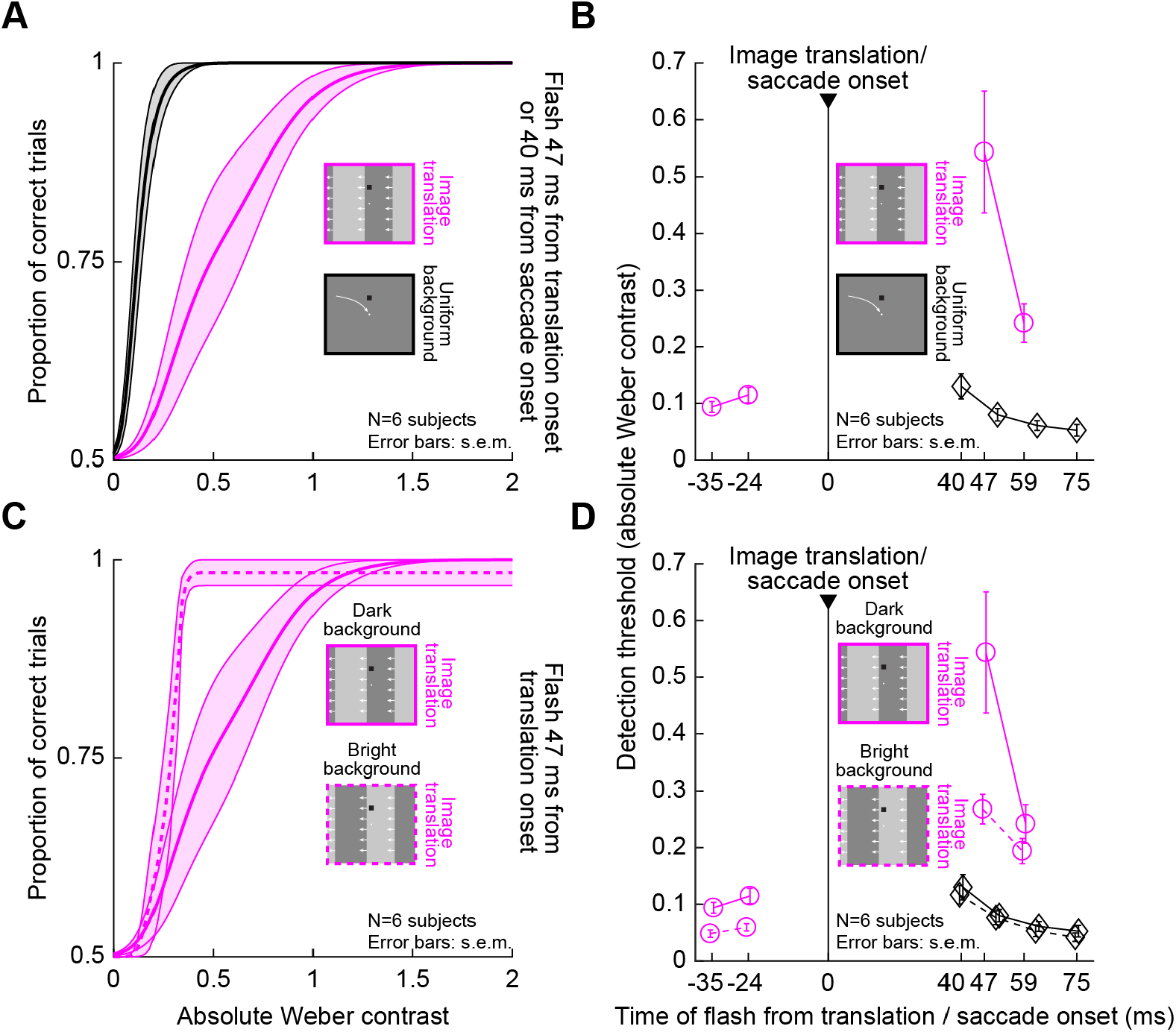
Highly similar visual dependencies of perceptual suppression in the absence of saccades. **(A)** Psychometric curve of perceptual detectability from condition 4, in which a saccade-like displacement of a vertical luminance edge took place (conceptually similar to condition 3 with real saccades). For reference, the black curve shows the psychometric curve from condition 1 (real saccades over a uniform background) at 40 ms. The saccade-like image displacement caused large suppression (a larger rightward shift of the curve), similar to the large suppression seen with real saccades (e.g. Fig. 4). Error bands denote s.e.m. across subjects. **(B)** Time course of perceptual suppression in condition 4. For reference, the time course of saccadic suppression over a uniform background (condition 1) is also shown in black. **(C)** There was also stronger suppression with (negative polarity) flashes over a dark background than with the same flashes over a bright background, exactly like with real saccades (e.g. Fig. 5). The average psychometric curve was shifted more to the right with a dark background (solid). **(D)** The dependence of perceptual suppression on background luminance was also clear in the whole time course, and this dependence was similar to that seen with real saccades in a similar visual condition of crossing an edge (e.g. Fig. 5). For reference, the black curves show the results obtained with real saccades across a uniform dark (solid) or bright (dashed) background (same as in Fig. 6).

Second, we confirmed that there was stronger perceptual suppression over a dark background versus over a bright background with our simulated saccades of condition 4. This effect can be seen in Fig. 7C, in which we plotted the average psychometric curves across all subjects at the same flash time, but with flashes occurring over either a bright or dark background. As we did above for simplicity in several other figures, we only plotted the curves for negative polarity flashes (but see below for explicit analysis of flash polarity in this condition as well). As can be seen, the psychometric curve for dark backgrounds (replicated from Fig. 7A) was shifted farther to the right compared to the curve for bright backgrounds, consistent with stronger perceptual suppression. In Fig. 7D, the full time courses can be seen for both background luminances. For reference, the curves for saccadic suppression with real saccades over a uniform background are also shown in black (condition 1). At the first time point after visual transient onset in the simulated saccade condition, threshold contrast was 0.54 +/- 0.11 s.e.m. and 0.27 +/- 0.03 s.e.m. Weber contrast for dark and bright backgrounds, respectively (paired t-test: t(5) = 4.5, p = 0.006).

Finally, we also confirmed that the same interactions between flash polarity and background luminance also persisted with simulated saccade-like image shifts. Specifically, we used condition 4 to explore psychometric curves of perceptual detectability when flashes appeared at different times relative to image displacement onset and on different background luminances. Figure 8A shows such curves for the case of negative or positive polarity flashes appearing over a dark background 47 ms after the onset of rapid image displacement. The curve for the negative polarity flashes was shifted more to the right than the curve for positive polarity flashes, suggesting stronger perceptual suppression. This is similar to what we also saw with real saccades (Fig. 6). Indeed, the time courses (Fig. 8B, C) exhibited very similar dependencies on flash polarity to the real saccade conditions. For example, over a dark background (Fig. 8B), perceptual thresholds were at 0.54 +/- 0.11 s.e.m. Weber contrast for negative polarity flashes, and 0.35 +/- 0.04 s.e.m. for positive polarity flashes, at the time closest to peak suppression in our data (47 ms). This was statistically significant (paired t-test: t(5) = 2.7, p = 0.04). For reference, Fig. 8B also shows the thresholds from condition 1 with real saccades over a uniform background. As can be seen, the suppression was significantly stronger with translation of a luminance edge across the retina than with real saccades over a uniform background. In fact, the threshold values in condition 4 (Figs. 7, 8) immediately after simulated saccade onset were generally similar in strength to those obtained immediately after real saccades in condition 3 (e.g. Figs. 4, 5); however, we caution against direct quantitative comparison, especially given the very different time courses of real versus simulated saccadic suppression profiles that can exist (Idrees, Baumann, Franke, et al., 2020).

**Figure 8.**
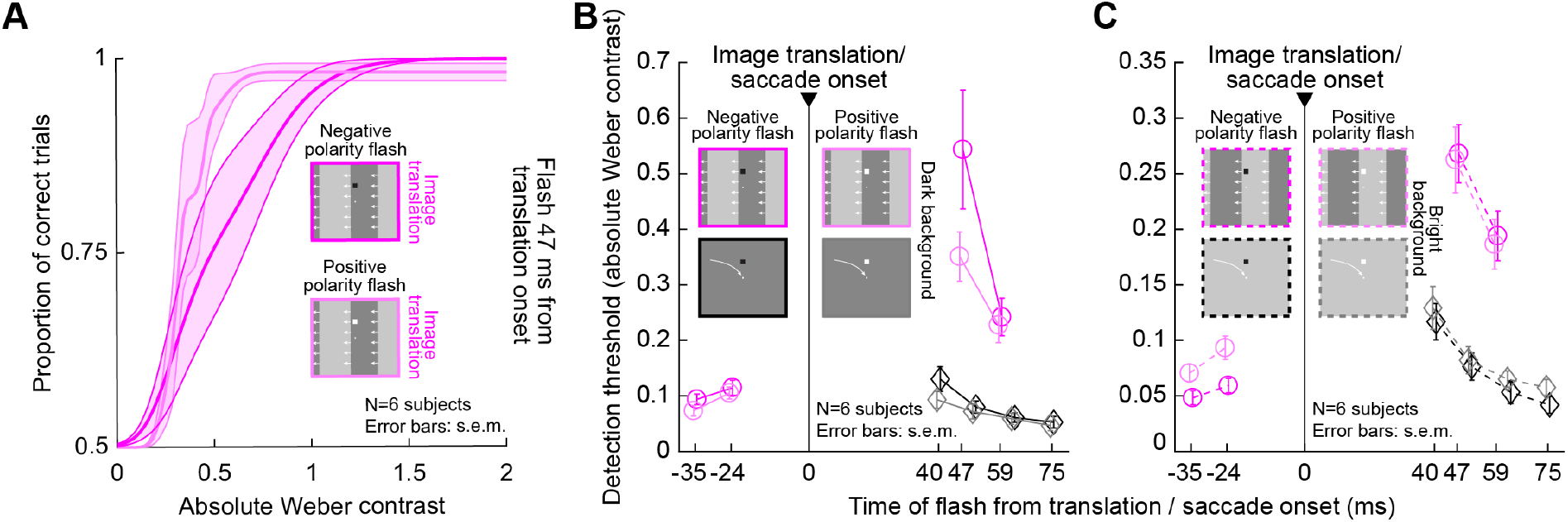
Stronger suppression for negative polarity flashes over dark backgrounds when compared to positive polarity flashes, even in the absence of saccades. **(A)** Average psychometric curves for negative and positive polarity flashes occurring at 47 ms after image translation onset, and appearing over a dark background. Saturated colors show results with negative polarity flashes, and unsaturated colors show results with positive polarity flashes. Error bands denote s.e.m. across subjects. Negative polarity flashes were associated with stronger perceptual suppression than positive polarity flashes. **(B)** Perceptual thresholds across subjects as a function of flash time after image translation onset in condition 4, comparing negative and positive polarity flashes over a dark background (error bars denote s.e.m. across subjects). For reference, the time course of suppression from condition 1 is also shown in black. Consistent with Fig. 7, perceptual suppression with an image translation (condition 4) was much stronger (higher perceptual thresholds) than in condition 1 with real saccades (compare colored to black curves). Moreover, perceptual saccadic suppression was stronger with negative polarity flashes than positive polarity flashes, especially near the time of peak suppression (47 ms). This is identical to the effect that we saw with real saccades in Fig. 6. **(C)** With bright backgrounds, the effects of flash polarity were absent, again consistent with what we observed with real saccades (Fig. 6).

The interaction between flash polarity and background luminance also extended to bright backgrounds. Specifically, with real saccades in conditions 2 and 3, we saw above (Fig. 6) that flash polarity did not alter saccadic suppression strength when flashes appeared over a bright background, unlike the case with flashes occurring over a dark background. This happened in a similar way in condition 4 with simulated saccades (Fig. 8C). Therefore, all of the visual dependencies that we observed with real saccades (Figs. 3-6) also occurred in the absence of any saccades, when saccade-like image translations were introduced (Figs. 7, 8).

It should be noted here that our simulated saccade condition was also important because it allowed us to additionally test probe flashes occurring before simulated saccade onset. In Figs. 7, 8, flashes with negative time were presented before the image translation. Nonetheless, they were still associated with threshold elevations, as we also recently reported in similar experiments (Idrees, Baumann, Franke, et al., 2020). For example, in Fig. 7B, perceptual threshold for a flash time of -24 ms was 0.11 Weber contrast, whereas it was only 0.09 at -35 ms. Therefore, there was elevation of threshold as time approached the onset of the image translation. These results, combined with earlier published work in the literature, make it likely that we would also observe pre-saccadic suppression in conditions 1, 2, and 3 above if we were to present flashes before the real saccades. Interestingly, even pre-translation flashes were clearly associated with stronger suppression when they occurred over a dark background as opposed to a bright background (Fig. 7D), consistent with the post-translation flash times demonstrating an influence of background luminance, which are in turn consistent with real saccade background effects.

### Luminance steps without saccade-like image sweeps cause modest perceptual suppression

Finally, for completeness, we also tested simple luminance steps during fixation, with no image translations (condition 5; Methods). In this case, we previously found that perceptual suppression does occur (Idrees, Baumann, Franke, et al., 2020), although in that previous study, we did not explore certain factors, like flash polarity, in much detail.

With luminance steps, perceptual suppression was significantly weaker in strength than with an image translations. For example, Fig. 9A shows thresholds when a flash occurred over a dark background, and Fig. 9B shows thresholds when a flash occurred over a bright background. For reference, saccadic suppression thresholds from condition 1 (i.e. with saccades across a uniform background) are also shown in black. Both conditions had the weakest overall suppression strengths in the whole study, and peak suppression in them was very similar. For example, at the first flash time after contrast change or saccade onset, the overall thresholds in both conditions were less than 0.18 Weber contrast (Fig. 9), which is much less than threshold contrasts in all other experiments with luminance edges or stripes (whether with or without saccades). This suggests an interesting role of luminance transients over peri-foveal image regions in perceptual saccadic suppression. Specifically, peri-foveally, the uniformity of the display was similar during the real saccades (condition 1) and the contrast change experiment (condition 5); both conditions were lacking local image translation across the retinal regions that were also probed with brief flash stimuli. Even though the entire retinal image (including display outer edges) were translated across the retina during real saccades, it seems that such a local image translation is necessary to maximize perceptual suppression of temporally-close probe flashes.

**Figure 9.**
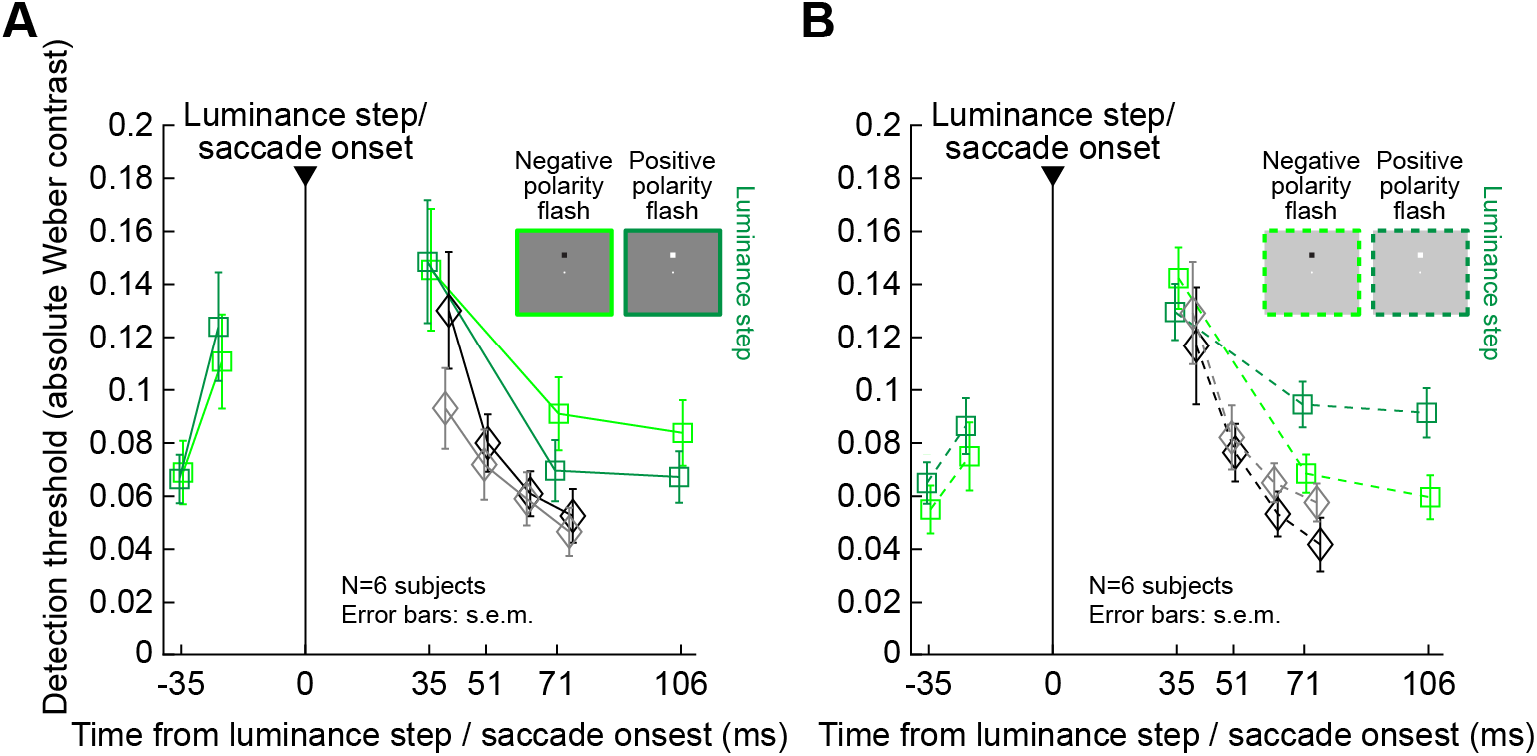
Weak perceptual suppression with simple luminance steps during fixation. **(A)** Perceptual contrast thresholds from condition 5 with just luminance steps (colored curves). Each curve shows results for either a positive (dark green) or negative (light green) polarity flash over a dark background, and error bars denote s.e.m. across subjects. For reference, thresholds from condition 1 are shown in black. Peak suppression with a luminance step was as weak as the suppression with real saccades over a uniform background (this was our weakest suppression condition in all experiments; Figs. 1-8). Moreover, during recovery, negative polarity flashes were harder to detect than positive polarity flashes. **(B)** Similar results for flashes occurring over a bright background. Now, it was positive polarity flashes that were harder to detect than negative polarity flashes during the recovery phase (at times 71 and 106 ms after luminance step occurrence). All other conventions are similar to Figs. 4, 5.

Figure 9 also reveals interesting differential effects of flash polarity on perceptual thresholds, but only at longer times after contrast change. Specifically, at both 71 ms and 106 ms after background luminance change (from bright to dark or vice versa), there was an interaction between flash polarity and background luminance: negative polarity flashes were harder to detect compared to positive polarity flashes over dark backgrounds, whereas positive polarity flashes over bright backgrounds were harder to detect than negative polarity flashes. To statistically validate this, we pooled the two latest flash times together and compared, with a paired t-test for each background, whether the detection thresholds differed for the two flash polarities. With a dark background, the positive polarity (mean threshold: 0.07 +/- 0.01 s.e.m. Weber contrast) significantly differed from the negative polarity (mean threshold: 0.09 +/- 0.01 s.e.m.) (t(5) = 5.8, p = 0.0022). Over a bright background, the positive polarity (threshold: 0.09 +/- 0.01 s.e.m. Weber contrast) significantly differed from the negative polarity (0.06 +/- 0.01 s.e.m.) (t(5) = -16, p <10^−4^). This effect was not present in any of our real or simulated saccade conditions (conditions 1-4), and it also might be a simple instantiation of Weber’s law, irrespective of perceptual suppression. Interestingly, in our recent investigation of a similar perceptual paradigm (Idrees, Baumann, Korympidou, et al., 2020), there was no clear impact of flash polarity. However, in that study, we did not use the more sensitive psychometric approach that we used here, so we could have been affected by floor and ceiling effects in that earlier study. Nonetheless, it is interesting that visual-visual interactions in saccadic suppression are best matched under fixation when saccade-like image displacements take place; other transients, such as just simple background luminance steps (Fig. 9), also cause mild suppression, but with different visual dependencies.

## Discussion

We investigated the visual components of perceptual saccadic suppression. We collected psychometric curves of perceptual detectability for brief peri-saccadic flashes and used these curves to estimate thresholds. We found that perceptual thresholds were only elevated mildly when saccades were made across a uniform background, but they were dramatically increased when gaze crossed a luminance edge (e.g. Figs. 3, 4). Moreover, the luminance at which gaze landed after saccade end did not have to be different to the luminance at saccade onset for the strength of saccadic suppression to increase; crossing a luminance stripe was also associated with very strong saccadic suppression when compared to the uniform background (condition, 2, e.g. Fig. 4). Interestingly, saccadic suppression was the strongest when the luminance of the background was dark, and there was an impact of flash polarity relative to the background luminance (particularly with negative polarity flashes over dark backgrounds; e.g. Figs. 5, 6). Critically, all of these visual dependencies of saccadic suppression still occurred when we replaced saccades with image displacements in the absence of any saccadic eye movements (e.g. Figs. 7, 8).

Our results from condition 3 confirm and extend those of (Maij et al., 2012). Specifically, in that study, the authors used a condition similar to ours (Fig. 1A, bottom), and they asked subjects to indicate whether they missed seeing a peri-saccadic flash or not (the overall context of the task was to study mislocalization of observed flash location, so the flashes were supra-threshold in general compared to ours). When the saccades were made over a uniform background, the subjects almost never missed the flashes. However, when the saccades involved gaze crossing a vertical luminance edge, there was an increased likelihood of misses. This is also what we saw with our more sensitive threshold measurements in the current study (e.g. Figs. 3, 4). As stated above, we also saw almost equally strong suppression when the edge was replaced with just a luminance stripe (condition 2, i.e. the starting and landing gaze positions were on the same background luminance; Figs. 3, 4). Combined, these results suggest that visual-visual interactions, in the form of gaze crossing a luminance pattern, matter a great deal for saccadic suppression.

Moreover, the effects are highly repeatable across individuals. In all analyses, average measurements across subjects were very consistent and repeatable, as evidenced by the small error bars in all our figures. Indeed, when we plotted the individual psychometric curves of each subject for one of the conditions, as in the example of Fig. 10, we saw remarkable repeatability across subjects. This was true in our earlier study as well, in which we saw high consistency across individual subjects (Idrees, Baumann, Franke, et al., 2020). Therefore, the effects that we report here are clearly robust across individuals.

**Figure 10.**
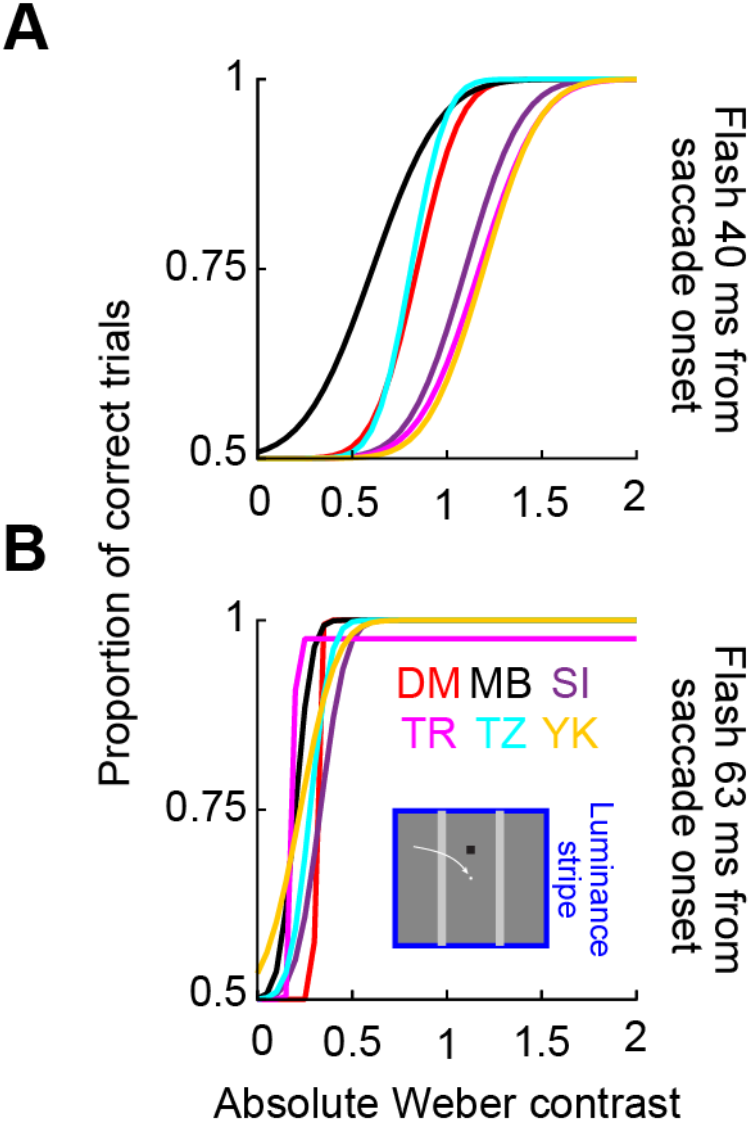
Raw psychometric curves at two flash times from each individual subject in condition 2. **(A)** Psychometric curves of all subjects with a negative polarity flash at 40 ms and dark backgrounds. **(B)** The same for the flash at 63 ms. There was strong coherence across subjects in the modulations of perceptual detectability in our experiments.

It is also interesting that the background luminance has a strong influence on the strength of saccadic suppression. In all analyses, we assessed perceptual thresholds based on calculating the absolute luminance difference between the probe flash and the background, normalized against the background luminance (i.e. Weber contrast). We observed that subjects needed more luminance contrast relative to the background (even with positive polarity flashes) to detect stimuli peri-saccadically (or peri-translation in condition 4) when the overall background was dark than when it was light. The fact that this effect was stronger with the dark background compared to the bright one rules out the possibility that this was a simple instantiation of basic Weber’s law (i.e. that more luminance needs to be added to a bright background than to a dark background for equal detectability). Rather, there was an apparent interaction between image translation and gaze finally encountering an image patch with a dark region. Similarly, there was also an interaction associated with the probe polarity itself: after saccades (or image translations), dark backgrounds required larger negative polarity contrasts for perceptual detection than positive polarity contrasts, but no polarity effects existed with a bright background (e.g. Fig. 6). It could be the case that crossing an image patch to a darker background induces a stronger visual transient (e.g. in the retina) than crossing an image patch to a brighter background, which then modifies the subsequent visual response to the probe flash itself. It would be interesting to explore this mechanism by relating how our observed interaction effects (that is, perceptually) relate to the activity of ON and OFF channels in ex-vivo mouse and pig retinae, similar to our recent investigations (Idrees, Baumann, Korympidou, et al., 2020). Intriguingly, preliminary data from that study indicate that ex-vivo non-human primate retinae may exhibit ON and OFF dependencies that might be similar to human effects (Idrees, Baumann, Korympidou, et al., 2020), at least in our condition 5. On the other hand, our probe flash polarity effects (e.g. Figs. 6, 8) demonstrate that strong perceptual suppression can occur irrespective of the combination of visual transient polarity and probe flash polarity; this suggests the presence of other suppressive mechanisms beyond the dynamic reversal suppression mechanism (which depends on a difference in polarity between visual transient and probe flash) that we uncovered in the mouse and pig retinae in the same study. Of course, it would also be interesting to investigate the visual-visual interactions of saccadic suppression in neural circuits downstream of the retina in non-human primates (or other relevant species used in systems neuroscience).

The importance of visual-visual interactions in perceptual saccadic suppression also makes us consider the possibility that with our uniform background condition (condition 1), the edges of the monitor may have had a non-negligible modulatory effect on the phenomenon. Specifically, for both the bright and dark background luminances used in the uniform background condition, the display monitor was still brighter than the rest of the laboratory. Therefore, there was a rectangular frame that was moved on the retina by the saccades. This likely affected perception since our other conditions (with luminance edges crossing the fovea) had a very strong effect on the strength of saccadic suppression. Nonetheless, it is still intriguing that, by far, the largest effects on peri-saccadic perceptual thresholds that we observed required that the luminance edges or patterns cross the fovea during the saccades. Such foveal crossing (or, more generally, crossing of the retinal patch experiencing the probe flashes) also happened in our simulated saccade condition (condition 4), which again showed strong perceptual suppression effects. This could be a function of the spatial resolution of the retinal regions (and associated downstream visual areas) that experience temporal luminance modulations during the saccades and the flashes.

In addition to the above effects in real saccades, we also saw essentially the same visual dependencies in experiments with visual-visual interactions no longer involving real saccades. Specifically, in our condition 4, we translated the luminance edge across the fovea and presented probe flashes near the time of the translation. We observed the same dependencies on background luminance, and the same interactions with flash luminance polarity, as with real saccades. This confirms our hypothesis (Idrees, Baumann, Franke, et al., 2020; Idrees, Baumann, Korympidou, et al., 2020) that visual-visual interactions play an important role in saccadic suppression, and it also extends the image conditions under which we could observe remarkable similarities between real and simulated saccades in terms of perceptual suppression (Idrees, Baumann, Franke, et al., 2020). All of this is in agreement with other phenomena, like the fact that intra-saccadic motion can still be perceived if its speed is within the detectability range of our motion sensors in the brain (Castet et al., 2001; Castet & Masson, 2000), and also the fact that intra-saccadic motion streaks provide important reference frames for keeping track of object locations (Schweitzer & Rolfs, 2020). This evidence highlights the role of vision even during rapid eye movements. In that sense, suppression occurring before saccades or image translations (e.g. Figs. 7-9) may be viewed as reflecting visual-visual interactions such as backwards masking (Breitmeyer, 2007; Brooks & Fuchs, 1975; Judge, Wurtz, & Richmond, 1980; Mackay, 1970; Mitrani et al., 1975).

Naturally, a role for prior knowledge of movement commands at the time of saccade generation (e.g. corollary discharge) must additionally matter for saccadic suppression, and the real question becomes how visual and non-visual mechanisms interact in this phenomenon. In other words, it need not be the case that saccadic suppression is only purely motor or only purely visual. For example, experiments under conditions of whiteout (i.e. absolutely uniform illumination across the entire retina) might suggest a role for a movement-related impact on perceptual saccadic suppression (Riggs & Manning, 1982). Interestingly, at the neural level, enhancement, rather than suppression, seems to take place in human LGN and V1 across saccades without visual stimulation (Sylvester, Haynes, & Rees, 2005; Sylvester & Rees, 2006). This suggests a very different role for knowledge of saccade commands than simply to actively reduce visual sensitivity. Similarly, we recently found that while the visual properties of perceptual suppression were highly similar with or without real saccades, there was a massive difference in the time course of suppression when real saccades were generated: perceptual suppression was much more short-lived when it was associated with real saccades than when it was triggered by visual-visual interactions in the absence of saccades (Idrees, Baumann, Franke, et al., 2020). This is very reassuring, in retrospect, because of the strong need to minimize saccade-induced disruptions in vision as much as possible. However, it also raises highly interesting and unanswered questions on what exactly the role of saccade-related movement commands is in modulating the properties of perceptual saccadic suppression. Could it be that corollary discharge is there to shorten saccadic suppression, rather than to cause it? This is a topic that we think will lead to very interesting new neurophysiological insights on the field of trans-saccadic perception.

## Conclusions

The phenomenon of perceptual saccadic suppression possesses a strong visual component, reflecting interactions between visual activation caused by saccade-induced image translations and visual activation caused by the brief probe flashes used to measure the sensitivity of the visual system around the time of saccades. This visual component of saccadic suppression motivates future neurophysiological, perceptual, and theoretical investigations on how this component interacts with internal knowledge of saccadic movement generation commands (i.e. efference copies or corollary discharge) for minimizing the disruptions to vision that can be caused by saccades.

## Acknowledgements

MPB and ZMH were funded by the Deutsche Forschungsgemeinschaft (DFG) Collaborative Research Center SFB 1233 on “Robust Vision” (project number: 276693517). SI and TAM were funded by DFG grant number MU3792/3-1.

